# Antibody Display of cell surface receptor Tetraspanin12 and SARS-CoV-2 spike protein

**DOI:** 10.1101/2021.05.29.446300

**Authors:** Fu-Lien Hsieh, Tao-Hsin Chang

**Affiliations:** Department of Molecular Biology and Genetics, Johns Hopkins University School of Medicine, Baltimore, MD 21205, United States; Howard Hughes Medical Institute, Johns Hopkins University School of Medicine, Baltimore, MD 21205, United States

**Keywords:** Antibody, display technology, Tetraspanin, SARS-CoV-2, retinal and infectious diseases

## Abstract

In previous work, Hsieh and Higgins presented a novel structure of antibodies identified from malaria-exposed individuals, in which the extracellular immunoglobulin (Ig)-like domain of leukocyte-associated immunoglobulin-like receptor 1 (LAIR1) is presented on the third complementarity determining regions (CDR3) of the Ig heavy chain. Here we develop an Antibody Display technology based on this LAIR1-containing antibody, by grafting proteins of interest (POI) onto the heavy chain CDR3 while retaining the biological properties of the POI. As a proof of principle, we displayed the second extracellular domain of Tetraspanin12 (Tspan12_EC2_) and the receptor-binding domain (RBD) of SARS-CoV-2 spike protein on the heavy chain CDR3. Our data revealed that Antibody Display Tspan12_EC2_ bound to Norrie Disease Protein (Norrin) and Antibody Display SARS-CoV-2 RBD bound to angiotensin-converting enzyme 2 (ACE2) and neutralizing nanobodies. Collectively, Antibody Display technology offers the general strategy of designing novel antibodies by grafting POI onto the CDR3.

## Introduction

Antibodies are crucial reagents for biomedical research, diagnostics, and therapeutics. The conventional antibody, immunoglobulin G (IgG), has a highly conserved architecture that is composed of two heavy chains and two light chains (Owen et al., 2013). Each chain contains three hypervariable loops, known as complementarity-determining regions (CDRs), that form the antigen-binding site, which determines the binding specificity for antigens (Chothia et al., 1989). Unexpectedly, a group of unusual antibodies were identified recently from malaria-endemic regions of Africa (Tan et al., 2016). These antibodies have a remarkable feature of the antigen-binding site in which an intact collagen-binding domain, adopting the immunoglobulin (Ig)-like fold, from leukocyte-associated immunoglobulin-like receptor 1 (LAIR1) was inserted into the CDR3 of the heavy chain (**Figure 1A**). The LAIR1 insert carrying multiple somatic mutations interacts with the repetitive interspersed families of polypeptides (RIFIN) that are expressed on the surface of *Plasmodium falciparum*-infected erythrocytes (Tan et al., 2016). A detailed structural analysis of this LAIR1-containing antibody (MGD21) has been reported (Hsieh and Higgins, 2017). Pieper et al., further found that malaria-exposed individuals had two additional types of LAIR1-containing antibodies in which the LAIR1 insert was located between the variable domain of the heavy chain (V_H_) and the constant domain of the heavy chain (C_H1_) or fused to the FC fragment (Pieper et al., 2017).

**Figure 1.**
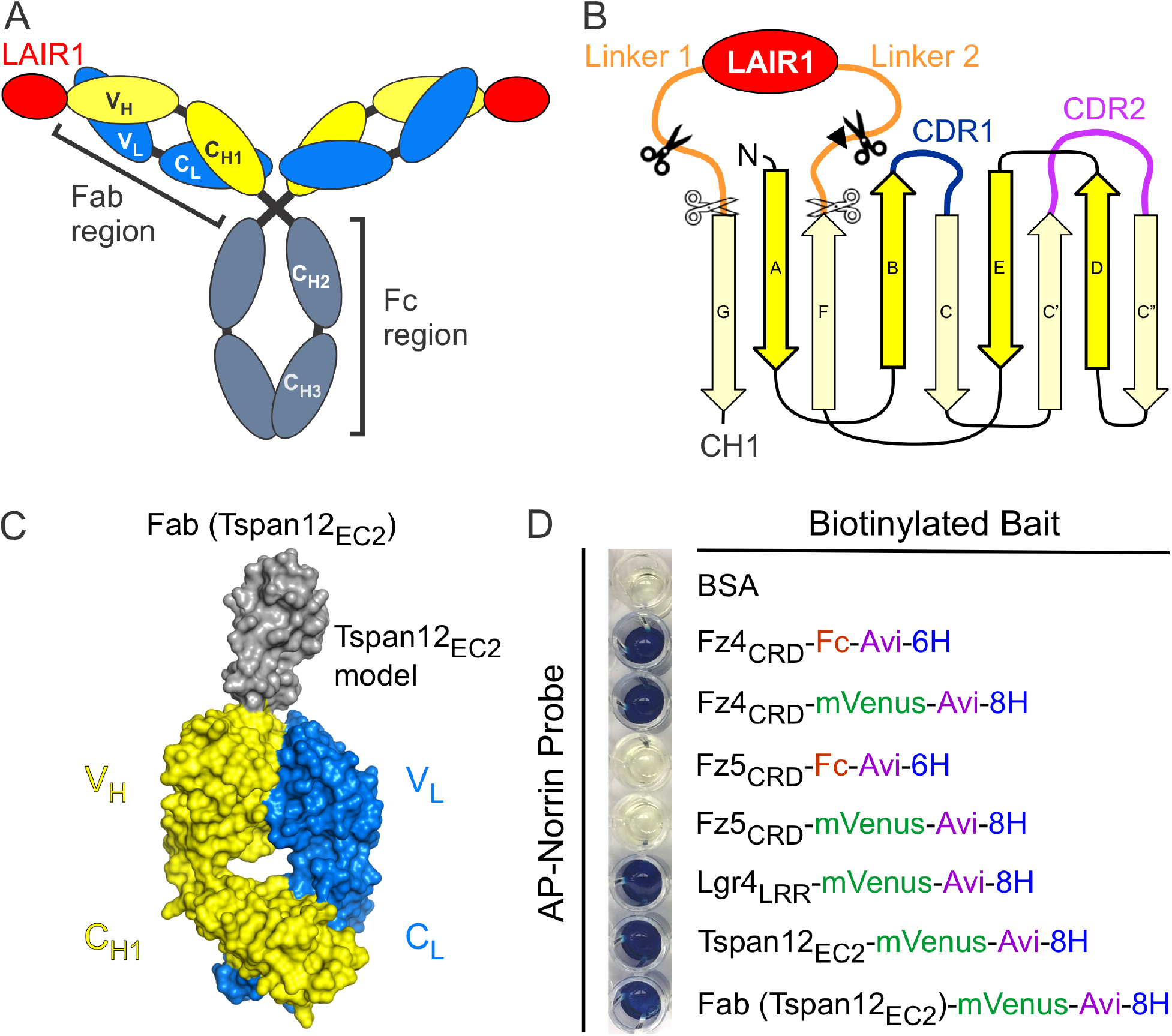
Antibody Display Tspan12_EC2_ (Tspan12_EC2_-AD). **(A)** Schematic representation of a LAIR1-containing antibody. Each Fab region is composed of variable domains of the heavy chain (V_H_) and the light chain (V_L_), constant domains of the heavy chain (C_H1_) and the light chain (C_L_), and a LAIRI insert on the V_H_. The Fc region contains constant domains of the heavy chain (C_H2_ and C_H3_). **(B)** Diagram representation of a LAIRI insert on the V_H_ and the construct design for Tspan12_EC2_-AD, see also **Figure 1–Figure supplement 2**. Tspan12_EC2_ replaced the LAIR1 insert (red) with two connecting linkers (orange) at different insertion sites (scissors) on the V_H_ (yellow). A black triangle indicates the location of C223 of the V_H_ forming a disulphide bond with C93 of the V_L_ (Notably, V_L_ C93A mutant was used, while the V_H_ construct without C223), see also **Figure 1–Figure supplement 2**. **(C)** Molecular model of Tspan12_EC2_-AD showing in the Fab format. **(D)** AP based protein-protein interaction assay. AP-Norrin (probe) incubated with biotinylated baits which were immobilized on streptavidin-coated wells. After washing unbound AP probes, the bound AP probes were visualized with BluePhos phosphatase substrate solution (a colorimetric AP reaction). BSA and Fz5_CRD_ fusion proteins are used as negative controls.

The Tetraspanin (Tspan) receptor family contains 33 members in humans and play central roles in diverse biological processes (cell-cell fusion, immune response, and vascularization) and diseases (pathogen infection and tumorigenesis) (Charrin et al., 2014; Hemler, 2014; Vences-Catalan and Levy, 2018). Previous structural analyses revealed that Tspan receptors adopt a conserved architecture composed of four transmembrane helices (TM1–TM4), intracellular N- and C-termini, a first and smaller extracellular domain (EC1; 13-30 residues) between TM1 and TM2, and a second and larger extracellular domain (EC2; 70-130 residues) between TM3 and TM4 (Kitadokoro et al., 2001; Oosterheert et al., 2020; Umeda et al., 2020; Yang et al., 2020b; Zimmerman et al., 2016). Functional studies showed that EC2 of Tspan receptors is typically a critical domain for the interactions with partner proteins or pathogens on the cell surface (Green et al., 2011; Rajesh et al., 2012; Zhu et al., 2002). Tspan12 is important in central nervous system (CNS) blood vessel development and blood-brain/retina barrier formation (Junge et al., 2009; Lai et al., 2017; Wang et al., 2018; Zhang et al., 2018). Genetic studies revealed that (1) deficiency of Tspan12 results in familial exudative vitreoretinopathy (FEVR), a hereditary disorder charactered by abnormal development of the retinal vasculature, often leading to retinal detachment and vision loss (Junge et al., 2009; Nikopoulos et al., 2010; Poulter et al., 2010), and (2) Tspan12 is a co-activator of Norrie Disease Protein (also named Norrin) mediated β-catenin signaling (Junge et al., 2009; Luhmann et al., 2005; Stenman et al., 2008; Xu et al., 2004). Norrin, a secreted cystine-knot growth factor, activates β-catenin signaling by binding to the extracellular cystine-rich domain (CRD) of Frizzled4 (Fz4) and the N-terminal domains of low-density lipoprotein receptor-related protein 5/6 (Lrp5/6) (Chang et al., 2015; Ke et al., 2013; Smallwood et al., 2007; Xu et al., 2004). Deficiencies of Fz4 and Lrp5 also result in FEVR (Robitaille et al., 2002; Toomes et al., 2004). Recently, Lai et al. suggested that the EC2 of Tspan12 (Tspan12_EC2_) is a critical region for Norrin binding (Lai et al., 2017). However, whether Tspan12_EC2_ interacts with Norrin directly remains obscure because of the technical challenge of obtaining recombinant Tspan12_EC2_ without its conserved transmembrane helices (Junge et al., 2009; Lai et al., 2017).

The worldwide spread of coronavirus disease-2019 (COVID-19) causing by severe acute respiratory syndrome coronavirus 2 (SARS-CoV-2) has become a global pandemic (Zhou et al., 2020a; Zhu et al., 2020). The surface of the SARS-CoV-2 particle is decorated with the heavily glycosylated trimeric spike (S) protein protruding from the virus membrane envelope (Fung and Liu, 2019; Shang et al., 2020; Walls et al., 2020; Wang et al., 2020b; Watanabe et al., 2020; Wrapp et al., 2020b; Yao et al., 2020). Specifically, SARS-CoV-2 S protein contains the receptor-binding domain (RBD) which mediates the interaction with a virus entry receptor, angiotensin-converting enzyme 2 (ACE2). Therefore, SARS-CoV-2 S protein is an attractive target for therapeutic and vaccine development (Hoffmann et al., 2020; Walls et al., 2020; Zhou et al., 2020a; Zhou et al., 2020b). SARS-CoV-2 S protein is immunogenic and the major target of neutralizing antibodies from COVID-19 convalescent individuals (Barnes et al., 2020; Baum et al., 2020; Brouwer et al., 2020; Chi et al., 2020; Ju et al., 2020; Seydoux et al., 2020; Walls et al., 2020; Wang et al., 2020a; Wu et al., 2020b). Furthermore, Piccoli et al., showed that more than 90% of these neutralizing antibodies from COVID-19 convalescent plasma target the RBD of SARS-CoV-2 S protein for protective humoral responses (Piccoli et al., 2020). However, recent studies showed that mutations in the RBD present in the current circulating SARS-CoV-2 variants decrease the potency of neutralizing antibodies (Greaney et al., 2021; Starr et al., 2021; Weisblum et al., 2020). Therefore, understanding how the RBD of SARS-CoV-2 S protein responds to ACE2 and neutralizing antibodies is critical for ongoing vaccine development, immunotherapy, and the assessment of the RBD mutations occurring in circulating SARS-CoV-2 stains.

In this report, we have developed an antibody-based display approach, termed Antibody Display, by grafting proteins of interest (POI) onto the heavy chain CDR3 to generate biological active and stably folded chimera antibodies. As a proof of principle, we used the structure-guided protein design approach based on the structure of LAIR1-containing antibody (Hsieh and Higgins, 2017) to display the EC2 of Tspan12 and the RBD of SARS-CoV-2 S protein as insertions at the tip of the heavy chain CDR3. We showed that Antibody Display Tspan12_EC2_ (Tspan12_EC2_-AD) bound to Norrin and Antibody Display SARS-CoV-2 RBD (RBD-AD) bound to ACE2 and two neutralizing nanobodies (VHH-72 and H11-D4). We also designed a humanized and engineering variable domain of the heavy chain alone to display the RBD of SARS-CoV-2 and confirmed its binding properties.

## Result

### Design, expression and purification of Antibody Display Tspan12_EC2_

The design of the Antibody Display was informed by structural studies of LAIR1-containing antibody (**Figure 1A**) (Hsieh and Higgins, 2017). We hypothesized that POI could graft onto the heavy chain CDR3 via two short peptide linkers to generate chimeric antibodies while retaining the biological features of the POI. To test this concept, we focused initially on Tspan12_EC2_, because (1) Tspan12_EC2_ protrudes from two transmembrane helices of Tspan12 suggesting that the N- and C-termini of Tspan12_EC2_ could be linked to β-strands G and F of the heavy chain in a manner that preserves their native spacing (**Figure 1B**) and (2) the biomedical importance of Tspan12 in retinal vascular diseases.

We began our design by searching for homologous structures and conducting a multiple sequence alignment of Tspan25 (CD53) (Yang et al., 2020b), Tspan28 (CD81) (Kitadokoro et al., 2001; Zimmerman et al., 2016), and Tspan29 (CD9) (Oosterheert et al., 2020; Umeda et al., 2020) with Tspan12, as shown in **Figure 1–Figure supplement 1A**. The obtained information led us to generate constructs having different boundaries of Tspan12_EC2_ for expression trials in HEK293T cells (**Figure 1–Figure supplement 1B**). The constructs contained an N-terminal signal peptide and C-terminal monomeric Venus (mVenus) and 12xHis tags. To assess the level of secreted mVenus fusion protein expression, we used a time- and cost-efficient method, immobilized metal affinity chromatography (IMAC) followed by in-gel fluorescent imaging (Chang et al., 2020). Variations in the location of the Tspan12_EC2_ N- and C-termini produced large effects on protein yield (**Figure 1–Figure supplement 1C**). Construct 4 (Tspan12 residues 116-220) exhibited the highest yield, somewhat less than 0.1 mg/L, consistent with previous studies showing the technical difficulties in producing recombinant Tspan12_EC2_ (Junge et al., 2009; Lai et al., 2017). As the secretory pathway in mammalian cells has stringent protein quality-control machinery to ensure that secreted proteins are folded correctly (Trombetta and Parodi, 2003), the expression trials imply that residues 116-220 corresponds to a core folding unit of Tspan12_EC2_.

We next generated a three-dimensional model of Tspan12_EC2_ and designed Antibody Display Tspan12_EC2_ (Tspan12_EC2_-AD) model in a Fab format, Fab (Tspan12_EC2_), by computationally grafting the Tspan12_EC2_ onto the heavy chain CDR3 based on the structure of LAIR1-containing antibody (**Figure 1C**). We designed constructs based on these principles: (1) placing the core folding unit of Tspan12_EC2_ at the tip of the heavy chain CDR3 to maximally expose its binding regions for ligand binding, (2) preventing steric clashes between the Tspan12_EC2_ and the CDR loops of the heavy chain and light chain, and (3) testing variations in the length and flexibility of the connecting linkers between the Tspan12_EC2_ and the heavy chain. Five designs of these chimeric heavy chain constructs (**Figure 1–Figure supplement 2A**) were co-transfected with light chain constructs in HEK293T cells to yield secreted Fab proteins and were assessed for expression level using IMAC followed by in-gel fluorescent imaging (**Figure 1–Figure supplement 2B**). Interestingly, only two of these designs (construct 1 and 5) could be expressed as secreted Fab proteins by passing the protein quality-control machinery in mammalian cells (**Figure 1–Figure supplement 2B**). Specifically, Tspan12_EC2_-AD (construct 5) with the highest yield contained Tspan12 residues 115-220 and had the shortest connecting linkers (**Figure 1– Figure supplement 2B**). This Tspan12_EC2_-AD was produced at 1 mg/L, a yield close to that of the LAIR1-containing antibody (Hsieh and Higgins, 2017), and showed a 10-fold higher expression level than Tspan12_EC2_ (**Figure 1–Figure supplement 1C**).

### Functional analyses of Antibody Display Tspan12_EC2_

Next, to assess if Tspan12_EC2_-AD adopted the correct fold of Tspan12_EC2_, we tested the ability of recombinant Tspan12_EC2_ to bind to Norrin as measured by a colorimetric alkaline phosphatase (AP) reaction-based protein-protein interaction assay using an AP-Norrin fusion protein as a probe (Xu et al., 2004). Each bait protein had a biotinylated C-terminal Avi-tag and was captured on a streptavidin coated micro-well. Tspan12_EC2_ bound to AP-Norrin (**Figure 1D** and **Figure 1– Figure supplement 3**), as did the CRD of Fz4 (Fz4_CRD_) and the leucine-rich repeat domain of Leucine-rich repeat containing G-protein coupled receptor 4 (Lgr4_LRR_), extracellular domains that are known to bind to Norrin (Chang et al., 2015; Deng et al., 2013; Shen et al., 2015; Xu et al., 2004). As expected, AP-Norrin did not bind to Fz5_CRD_ (**Figure 1D**) (Chang et al., 2015; Smallwood et al., 2007). Importantly, Tspan12_EC2_-AD bound to AP-Norrin as did Tspan12_EC2_ (**Figure 1D**). Taken together, these results show that (1) EC2 of Tspan12 mediates Norrin binding, in agreement with previous genetic studies using domain swaps within full-length Tspan12 together with co-immunoprecipitation (Lai et al., 2017), and (2) correctly folded Tspan12_EC2_ can be grafted onto the heavy chain CDR3 without losing Norrin binding.

### Design, expression and purification of Antibody Display SARS-CoV-2 RBD

To assess whether the design principle of the Antibody Display can apply to other protein folds, we tested the RBD of SARS-CoV-2 S protein. We began the design of the Antibody Display SARS-CoV-2 RBD (RBD-AD) by searching the three-dimensional RBD structures of SARS-CoV-2 S protein (Barnes et al., 2020; Huo et al., 2020; Lan et al., 2020; Shang et al., 2020; Walls et al., 2020; Wang et al., 2020b; Wrapp et al., 2020b; Yuan et al., 2020; Zhou et al., 2020b). As shown in **Figure 2A**, we computationally grafted the RBD structure of SARS-CoV-2 S protein onto the heavy chain CDR3 of LAIR1-containing antibody to generate a model in in a Fab format, Fab (RBD), by using the same design principles as Tspan12_EC2_-AD.

**Figure 2.**
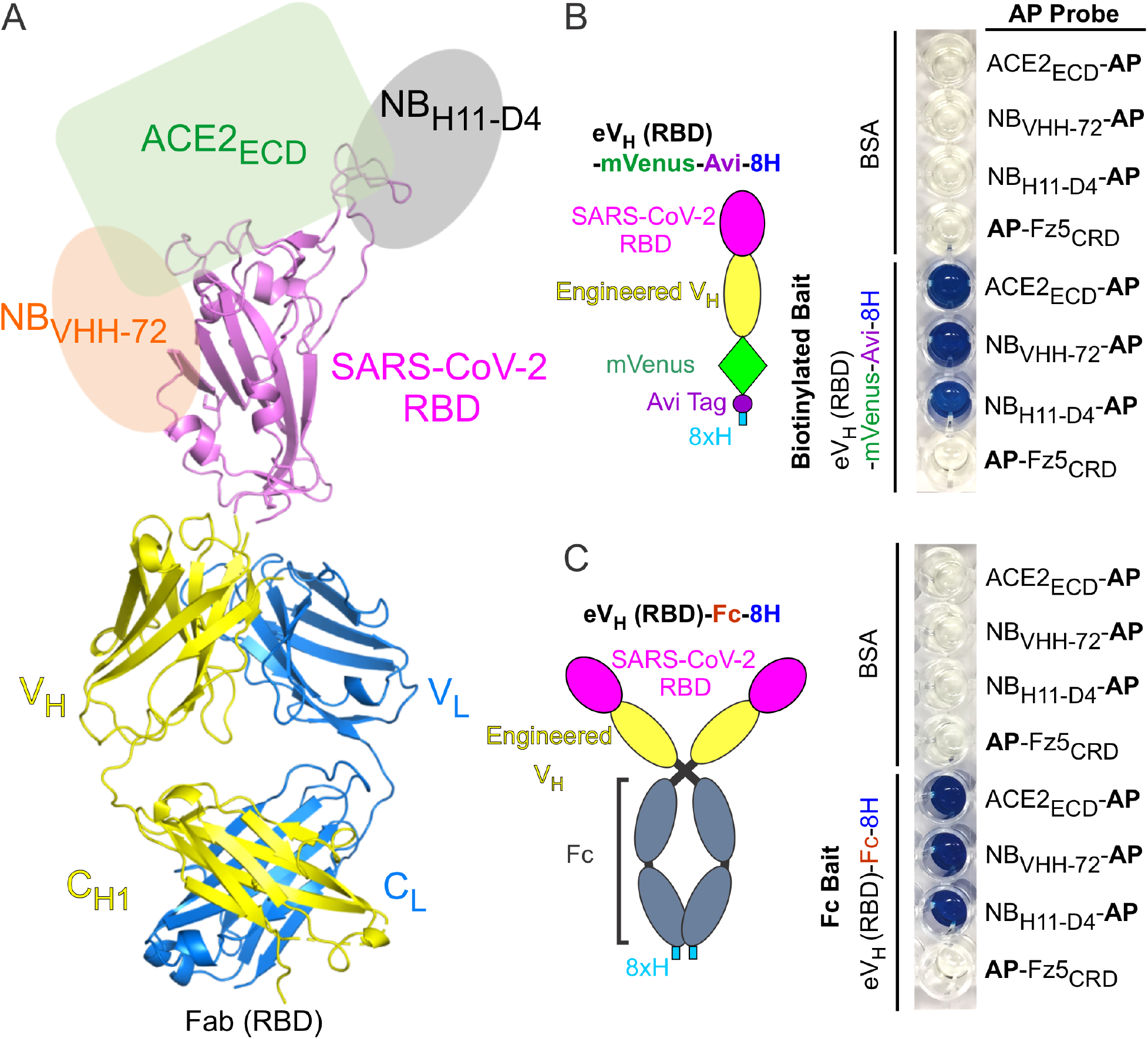
Antibody Display SARS-CoV-2 RBD (RBD-AD). **(A)** Molecular model of RBD-AD showing in the Fab format. The RBD (magenta) of SARS-CoV-2 S protein is displayed on the V_H_ (yellow). A schematic diagram shows the binding sites on the RBD for ACE2_ECD_ (green), NB_VHH-72_ (orange), and NB_H11-D4_ (grey). **(B)** A diagram of the RBD of SARS-CoV-2 S protein is displayed on the engineered V_H_ fused to a mVenus, an Avi tag, and an 8xH tag at the C-terminus. For the AP based binding assay, biotinylated baits were immobilized on streptavidin-coated wells. Bound AP fusion proteins (probes) were visualized with BluePhos phosphatase substrate solution. BSA and Fz5_CRD_ fusion proteins are used as negative controls. **(C)** Diagram of SARS-CoV-2 RBD-containing engineered V_H_ fused an 8xH tagged human Fc. FC baits were captured on protein-G coated wells and incubated with AP fusion proteins (probes). The bait and probe interactions were detected as in panel (B).

Furthermore, we hypothesized that a single V_H_ domain could serve as a scaffold to display SARS-CoV-2 RBD in the absence of light chains. Indeed, human single V_H_ domain mimicking nanobodies from camels has been developed as a minimum size of antigen recognition unit in the absence of light chains (Davies and Riechmann, 1995; Hamers-Casterman et al., 1993), although specific hallmark mutations are required to maintain the solubility and the binding property of the single V_H_ domain (Davies and Riechmann, 1995; Hamers-Casterman et al., 1993; Muyldermans, 2013). More recently, Wu et al., identified a highly soluble and stable single V_H_ domain from the human germline Ig heavy chain variable region 3-66*1 (IGHV3-66*1) allele (Wu et al., 2020a). Therefore, we next designed an engineered single V_H_ domain (eV_H_) based on the IGHV3-66*1 for the display of SARS-CoV-2 RBD, eV_H_(RBD), as shown in **Figure 2– Figure supplement 2**.

### Functional analyses of Antibody Display SARS-CoV-2 RBD

To verify whether RBD-AD retained binding to ACE2 and neutralizing antibodies, we generated AP fusion proteins with the extracellular domain of ACE2 (ACE2_ECD_-AP) and two anti-RBD nanobodies (NB_VHH-72_-AP and NB_H11-D4_-AP) (Huo et al., 2020; Wrapp et al., 2020a) (**Figure 2– Figure supplement 2**). We next prepared bait proteins of RBD-AD: (1) Fab (RBD) and eV_H_(RBD) constructs had a C-terminal mVenus and biotinylated Avi-tag, immobilized on streptavidin-coated wells (**Figure 2B** and **Figure 2–Figure supplement 3**), and (2) an eV_H_(RBD) construct contained the human IgG Fc fusion at the C-terminus, immobilized on a protein G-coated well (**Figure 2C**). As measured the binding by AP reaction-based assays, Fab (RBD) and eV_H_(RBD) bound robustly and specifically to ACE2_ECD_-AP, NB_VHH-72_-AP, and NB_H11-D4_-AP (**Figure 2B-C** and **Figure 2–Figure supplement 3**). We conclude that RBD of SARS-CoV-2 S protein can be displayed on the heavy chain CDR3 in functional form.

## Discussion

We present a novel antibody engineering approach, named Antibody Display, by grafting POI onto the heavy chain CDR3, which were designed through a structure-guided approach built on the architecture of LAIR1-containing antibodies naturally occurred in malaria-exposed individuals (Hsieh and Higgins, 2017; Tan et al., 2016). Most importantly, each POI retained its biological properties by taking advantage of the structural conservation and stability of the antibody Ig fold. As a proof of principle, we showed that the design and production of Tspan12_EC2_-AD and RBD-AD were successful. Colorimetric AP reaction-based binding assays further revealed that Tspan12_EC2_-AD bound to Norrin and RBD-AD bound to ACE2 and NBs (VHH-72 and H11-D4). Our findings suggest a generic design principle of the Antibody Display, particularly for the POI adopting a non-Ig like fold: (1) a known structure or a computational model derived from homologous protein structures, (2) N and C termini in proximity to graft onto the β-strands G and F of the heavy chain, and (3) prevention of steric clashes with the antibody. At present, the limitations for selecting POI for Antibody Display such as protein shape and size remain unclear; further studies will be required to address these questions.

Comparing with conventional Fc fusion proteins in which the POI is fused to the N terminus of Fc usually including the flexible hinge region, we reasoned that the Antibody Display, which grafts the POI onto the β-strands G and F of the heavy chain, provides several potential advantages: (1) design and production of a more conformationally constrained chimeric protein, (2) reduction of proteolysis with the POI inserted into a stable antibody Ig fold, and (3) extension of the current antibody engineering toolkit for generating bispecific and chimeric antibodies. Further studies will be needed to explore these potential advantages.

An intriguing feature of using the Antibody Display for Tspan12_EC2_ is with a better yield of protein production than Tspan12_EC2_. Our functional assays demonstrated that Tspan12_EC2_ bound to Norrin directly, in agreement with previous genetic studies (Lai et al., 2017). As Tspan12 playing critical roles in CNS vascular development (Junge et al., 2009; Zhang et al., 2018) and tumorigenesis (Knoblich et al., 2014; Otomo et al., 2014), Tspan12_EC2_-AD may be a useful reagent for these studies. Moreover, Tspan proteins are well known for their important roles in regulating tumor migration, invasion and metastasis (Charrin et al., 2014; Hemler, 2014; Vences-Catalan and Levy, 2018). Antibodies targeting Tspan29 (CD9), particularly on its EC2 region, have been shown to inhibit the progression of colorectal and gastric cancers (Nakamoto et al., 2009; Ovalle et al., 2007). Furthermore, Tspan proteins are used by several pathogens for infection (Charrin et al., 2014). For example, Hepatitis C Virus and *P. falciparum* use Tspan28 (CD81) for infection of hepatocytes (Pileri et al., 1998; Silvie et al., 2003). However, research on Tspan receptors has been hampered by the lack of an efficient approach to produce recombinant EC2 proteins. So far, only Tspan25 (CD53), Tspan28 (CD81), and Tspan29 (CD9) have been produced for biochemical and structural analyses among 33 Tspan receptors (Kitadokoro et al., 2001; Oosterheert et al., 2020; Umeda et al., 2020; Yang et al., 2020b; Zimmerman et al., 2016). More studies will be required to investigate whether the Antibody Display can apply to other Tspan proteins.

SARS-CoV-2 has resulted in a global pandemic and several types of COVID-19 vaccines such as nucleic acids, viral vectors, and protein subunits have been developed for vaccinations globally (Connors et al., 2021). Ongoing studies have shown that the RBD of SARS-CoV-2 S protein is the major target of neutralizing antibodies from COVID-19 convalescent individuals (Barnes et al., 2020; Baum et al., 2020; Brouwer et al., 2020; Chi et al., 2020; Ju et al., 2020; Piccoli et al., 2020; Seydoux et al., 2020; Walls et al., 2020; Wang et al., 2020a; Wu et al., 2020b), consistent with recent findings showing that the RBD, particularly decorated on nanoparticles, can induce a protective humoral immune response against SARS-CoV-2 infection (Cohen et al., 2021; Tan et al., 2021; Yang et al., 2020a). In this report, we demonstrated that the Antibody Display can be used to display the RBD of SARS-CoV-2 S protein on the heavy chain CDR3 without losing its binding properties. It is not yet known whether RBD-AD can induce neutralizing antibodies, but it would be an interesting test case and it could be relevant for the future preparedness and response capacity against emerging SARS-CoV-2 variants.

## Materials and Methods

### Construct design and molecular cloning

The human complementary DNA clones of Fz4, Fz5, and Tspan12 were gifts from Jeremy Nathans and the human ACE2 clone (Shang et al., 2020) was obtained from Addgene (code 145033). Synthetic DNA fragments of engineered V_H_ containing SARS-CoV-2 RBD (resi 333-527; with a P527A mutation; UniProtKB code P0DTC2; **Figure 2–Figure supplement 2**), NB_VHH-72_ (PDB code 6WAQ), and NB_H11-D4_ (PDB code 6YZ5) were obtained from Integrated DNA Technologies. Expression plasmids of MGD21 heavy chain pOPINVH and MGD21 light chain pOPINVL were described Hsieh and Higgins, 2017. The AP fusion Norrin construct was described in Xu et al., 2004.

The following five backbone constructs used for the Fc fusion or biotinylated baits and AP fusion probes were derived from the pHLsec vector (Aricescu et al., 2006). (1) The pHLsec-3C-Fc (C103A)-8H vector contains a C-terminal Human Rhinovirus (HRV)-3C protease cleavage site followed by the human immunoglobulin heavy constant gamma 1 (resi 102-330; having a C103A mutation to remove an unpaired cysteine; UniProtKB code P01857) and an 8xHis tag. (2) The pHLsec-Fc-Avi-6H vector contains the human immunoglobulin heavy constant gamma 1 (resi 101-330; UniProtKB code P01857) followed by an Avi tag that can be biotinylated by BirA ligases and a 6xHis tag. (3) The pHLsec-3C-mVenus-Avi-8H vector derived from pHLsec-mVenus-12H (Chang et al., 2015) was tagged with a C-terminally HRV-3C protease cleavage site followed by a mVenus fusion protein, an Avi tag and finally an 8xHis tag. (4) The pHL-N-AP-Myc-8H vector was constructed N-terminally with the human alkaline phosphatase (AP; resi 1-506; NCBI code NP_001623) followed by a Myc tag and finally a C-terminal 8xHis tag. (5) The pHLsec-C-Myc-AP-8H vector contains a C-terminal Myc tag followed by the human AP (resi 18-506; NCBI code NP_001623) and finally an 8xHis tag.

Tspan12 coding segments for the EC2 were PCR-amplified and cloned into the pHLsec-mVenus-12H (Chang et al., 2020; Chang et al., 2015). Tspan12_EC2_ inserts were grafted into a modified MGD21 heavy chain vector containing a C-terminal HRV-3C protease cleavage site followed by a mVenus fusion protein and finally an 8xHis tag (pHLsec-mMGD21-3C-mVenus-Avi-8H), by using three-fragments overlapping PCR experiments. The MGD21 light chain mutant (C91A and C93A) vector was generated by using a two-step overlapping PCR method. A DNA segment coding for CRD of Fz4 (resi 40-179) was constructed into the pHLsec-Fc-Avi-6H and pHLsec-3C-mVenus-Avi-8H. The CRD of Fz5 (resi 31-181) were cloned into the pHLsec-Fc-Avi-6H, pHLsec-3C-mVenus-Avi-8H, and pHL-N-AP-Myc-8H vectors. The Lgr4_LRR_ was constructed into the pHLsec-3C-mVenus-Avi-8H vector. SARS-CoV-2 RBD insert was grafted into the pHLsec-mMGD21-3C-mVenus-Avi-8H vector. An engineered V_H_ containing SARS-CoV-2 RBD was constructed into the pHLsec-3C-mVenus-Avi-8H and pHLsec-3C-Fc (C103A)-8H vectors. ACE2_ECD_ (resi 19-615), NB_VHH-72_, and NB_H11-D4_ were cloned into the pHLsec-AP-8H vector. All constructs were confirmed by sequencing.

### Computational modeling and structure-based design

The HHpred server (Soding et al., 2005; Zimmermann et al., 2018) was used to search for homologs of Tspan12_EC2_ in the protein structure database of the Protein Data Bank (PDB). The structures of Tspan28 (CD81; PDB codes 1G8Q and 5TCX) were selected to generate an initial computational model of Tspan12_EC2_ with Modeller (Eswar et al., 2006). Notably, the structures of Tspan25 (CD53; PDB code 6WVG), and Tspan29 (CD9, PDB codes 6RLR and 6K4J) were not available during the HHpred search. For the protein design of Antibody Display Tspan12_EC2_, LAIR1 insert was removed from the model of MGD21 Fab fragment (PDB code 5NST) and the Tspan12_EC2_ model was docked into the MGD21 V_H_ manually by using COOT (Emsley et al., 2010) and PyMOL Molecular Graphic System (Schrödinger, LLC). For the structure-guided design of Antibody Display SARS-CoV-2 RBD, the crystal structure of SARS-CoV-2 RBD (PBD code 6YLA) was selected and docked into the V_H_ region of the antibody manually by using COOT (Emsley et al., 2010).

### Protein expression and purification

HEK293T (ATCC CRL-11268) cells were maintained and transfected with the DNA using polyethylenimine (MilliporeSigma 408727) following the established procedures (Aricescu et al., 2006; Chang et al., 2015; Hsieh and Higgins, 2017; Hsieh et al., 2016). For Antibody Display Tspan12_EC2_, DNA constructs expressing heavy chain and light chain were mixed into a 1 to 1 ratio and co-expressed in HEK293T cells. For the biotinylated bait preparations, bait constructs were mixed with a pHLsec-BirA-ER vector (Chang et al., 2015) into a 3 to 1 ratio and co-transfected in HEK293T cells in the presence of 0.1 mM Biotin (MilliporeSigma B4639). The biotinylated baits were purified from conditioned media by the immobilized metal affinity chromatography (IMAC) method as previously described in (Chang et al., 2020). IMAC method was also used for the purification of His tagged Fc fusion and AP fusion proteins. To evaluate the expression level of secreted mVenus fusion proteins from conditioned media, IMAC followed by an in-gel fluorescence imaging method was used as previously described in (Chang et al., 2020).

### Alkaline Phosphatase based binding assay

The purified Fc fusion baits were captured on 96-well protein-G coated plates (Thermo Fisher Scientific 15131). The biotinylated baits were immobilized on 96-well streptavidin-coated plates (Thermo Fisher Scientific 15500). The wells were then washed three times with wash buffer (10 mM HEPES, pH 7.5, 0.15 M NaCl, 0.05% (w/v) Tween-20) and incubated with a 10-fold dilution of Blocker bovine serum albumin (BSA; Thermo Fisher Scientific 37525) in wash buffer for 1 h at 25 °C. The wells were washed with wash buffer and incubated with conditioned media containing AP probes at 4 °C overnight. The wells were subsequently washed three times with wash buffer and incubated with BluePhos phosphatase substrate solution (Kirkegaard and Perry Laboratories 50-88-00) to visualize the bound AP probes.

## Acknowledgements

We thank Jeremy Nathans for access to the laboratory and valuable comments on the manuscript; and Matthew Higgins for critical reading the manuscript and valuable comments. F-L H was supported by the Howard Hughes Medical Institute. T-H C was supported by a Human Frontier Science Program long-term fellowship (LT000130/2017-L).

## Contributions

F-L H and T-H C conceived the project, conducted experiments, and wrote the manuscript.

---

**Figure 1–Figure supplement 1.**
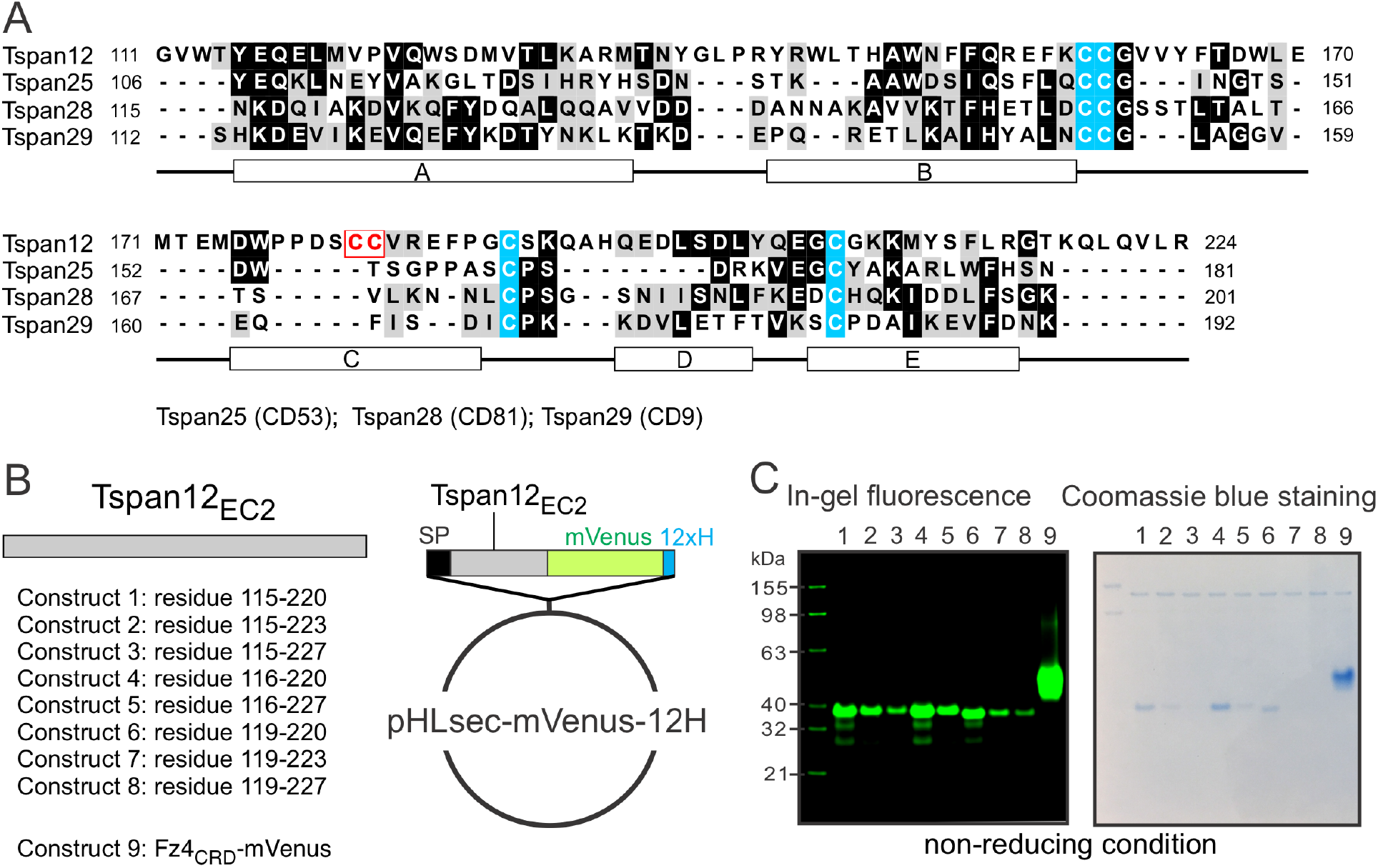
Tspan12_EC2_ sequence alignment, cloning, and expression trials. **(A)** Multiple sequence alignment of Tspan12_EC2_ with current available Tspan structures including Tspan25 (CD53; PDB code 6WVG), Tspan28 (CD81, PDB codes 1G8Q and 5TCX), and Tspan29 (CD9, PDB codes 6RLR and 6K4J). Identical residues are shaded in dark and similar residues in grey. Secondary structure elements (five α-helical regions A to E) are indicated based on the structures of Tspan25, Tspan28, and Tspan29. The conserved cysteine residues are highlighted in cyan. Tspan12 has two additional cysteine residues (a red box) conserved in the vertebrate. **(B)** Eight Tspan12_EC2_ expression constructs were cloned in the pHLsec-mVenus-12H vector. **(C)** Tspan12_EC2_ constructs having the mVenus fusion were transfected in HEK293T cells. Fz4_CRD_ with the mVenus fusion was used to compare the expression level. The conditioned media (1 ml) were purified by immobilized metal affinity chromatography (IMAC) and protein expression levels were determined by in-gel fluorescence imaging and Coomassie blue staining (Chang et al., 2020).

**Figure 1–Figure supplement 2.**
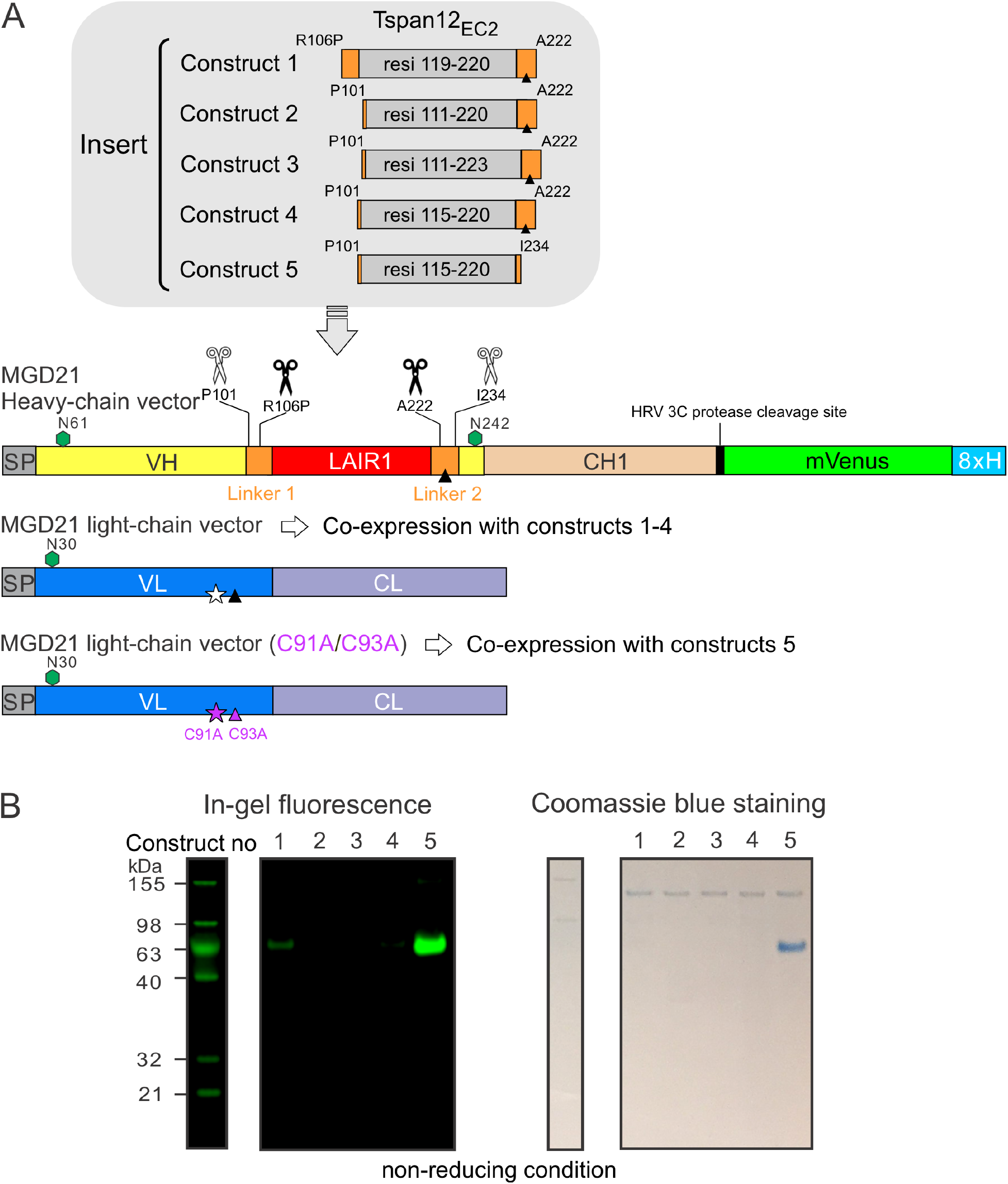
Construct design and protein expression of Tspan12_EC2_-AD. See also **Figure 1B** for a brief schematic representation of construct design. **(A)** Five inserts containing various fragments of Tspan12_EC2_ and connecting linker sequences were cloned into a modified antibody MGD21 heavy chain vector containing a HRV 3C protease cleavage site followed by a mVenus and an 8xHis tag. Constructs 1 to 4 were co-expressed with the MGD21 light chain vector in HEK293T cells. Construct 5 which lacks residue C223 on the V_H_ was co-transfected in HEK293T cells with an MGD21 light chain vector containing double mutations C91A and C93A (C93A mutation was used to block an inter-chain disulphide formation by V_H_ C223 and V_L_ C93; V_L_ C91 is an unpaired cysteine residue, so C91A mutation was designed to prevent the potential formation of artificial disulphide bonds). Green hexagons denote N-linked glycosylation sites. Black triangles show V_H_ C223. Star indicates V_L_ C91. **(B)** The conditioned media were harvested 2 days post-transfection and purified by IMAC. In-gel fluorescence imaging and Coomassie blue staining were used to evaluate protein expression levels.

**Figure 1–Figure supplement 3.**
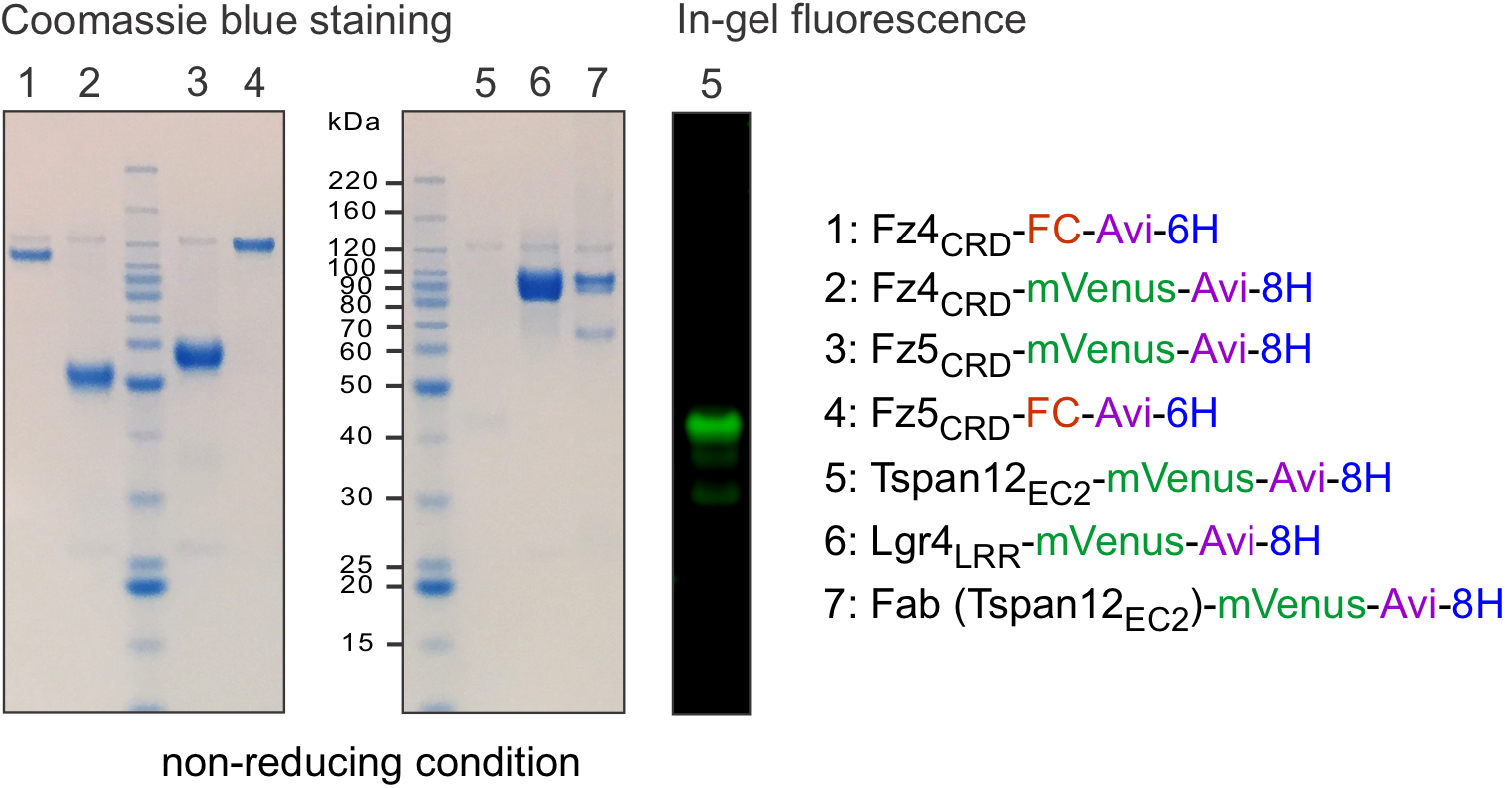
Coomassie blue-stained SDS-PAGE and in-gel fluorescence imaging showed the biotinylated baits that were purified by IMAC from the conditioned media and used for the AP based binding assay (**Figure 1D**).

**Figure 2–Figure supplement 1.**
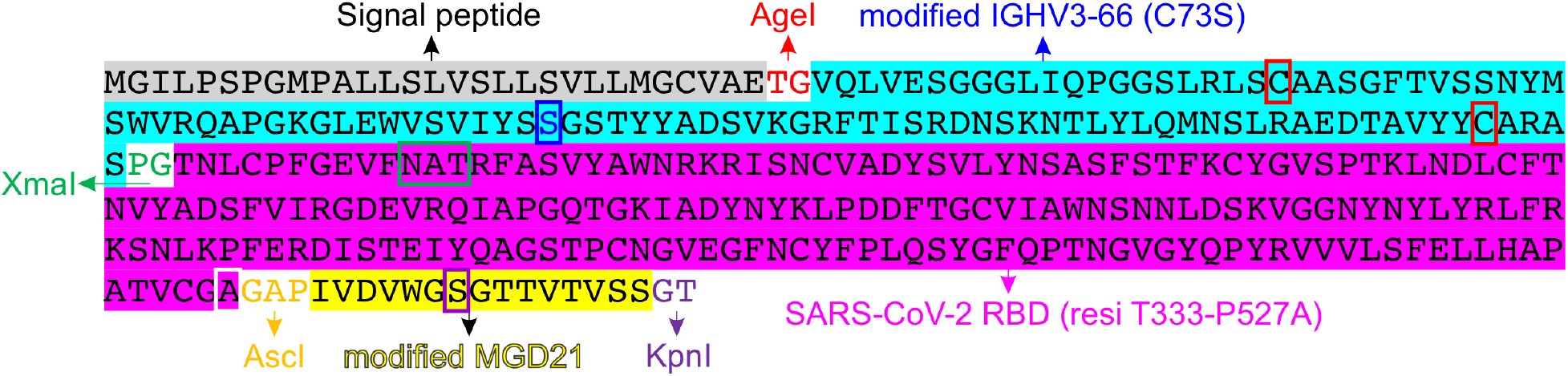
The amino acid sequence of the RBD of SARS-CoV-2 S protein displayed on the engineered V_H_. Engineered V_H_ is a chimeric fusion of the V_H_ region of IGHV3-66*1 highlighted in cyan (the C73S mutation is used to prevent the formation of artificial disulphide bonds and is boxed in blue) and a part of MGD21 V_H_ region highlighted in yellow (the N242S mutation is used to block the N-linked glycosylation and is boxed in purple; two residues, Y235 and Y236, are removed from the sequence by complementing the insertion of an AscI restriction site). The RBD (highlighted in magenta) of SARS-CoV-2 S protein contains a P527A mutation (boxed in white) by enhancing model restraint for the insertion. Red boxes denote two conserved cysteine residues of V_H_ forming a disulphide bond. A green box indicates an N-linked glycosylation site of the RBD of SARS-CoV-2 S protein.

**Figure 2–Figure supplement 2.**
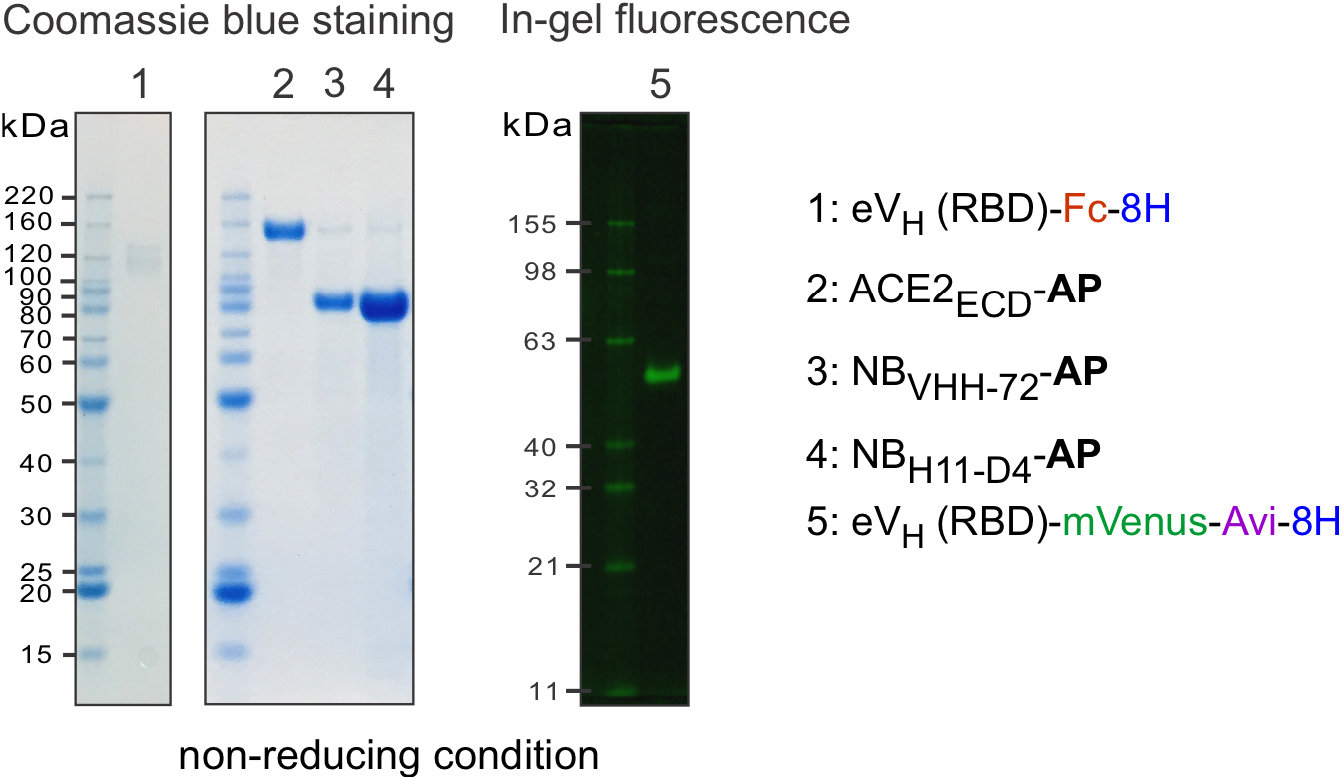
Coomassie blue-stained SDS-PAGE and the in-gel fluorescence results showed the biotinylated and Fc fusion baits and AP fusion probes that were purified by IMAC from the conditioned media. Constructs of ACE2_ECD_-AP, NB_VHH-72_-AP, and NB_H11-D4_-AP contain a C-terminal 8xHis tag. The AP based protein-protein interaction assays (**Figure 2B**, **Figure 2C** and **Figure 2–Figure supplement 3**) used the conditioned media.

**Figure 2–Figure supplement 3.**
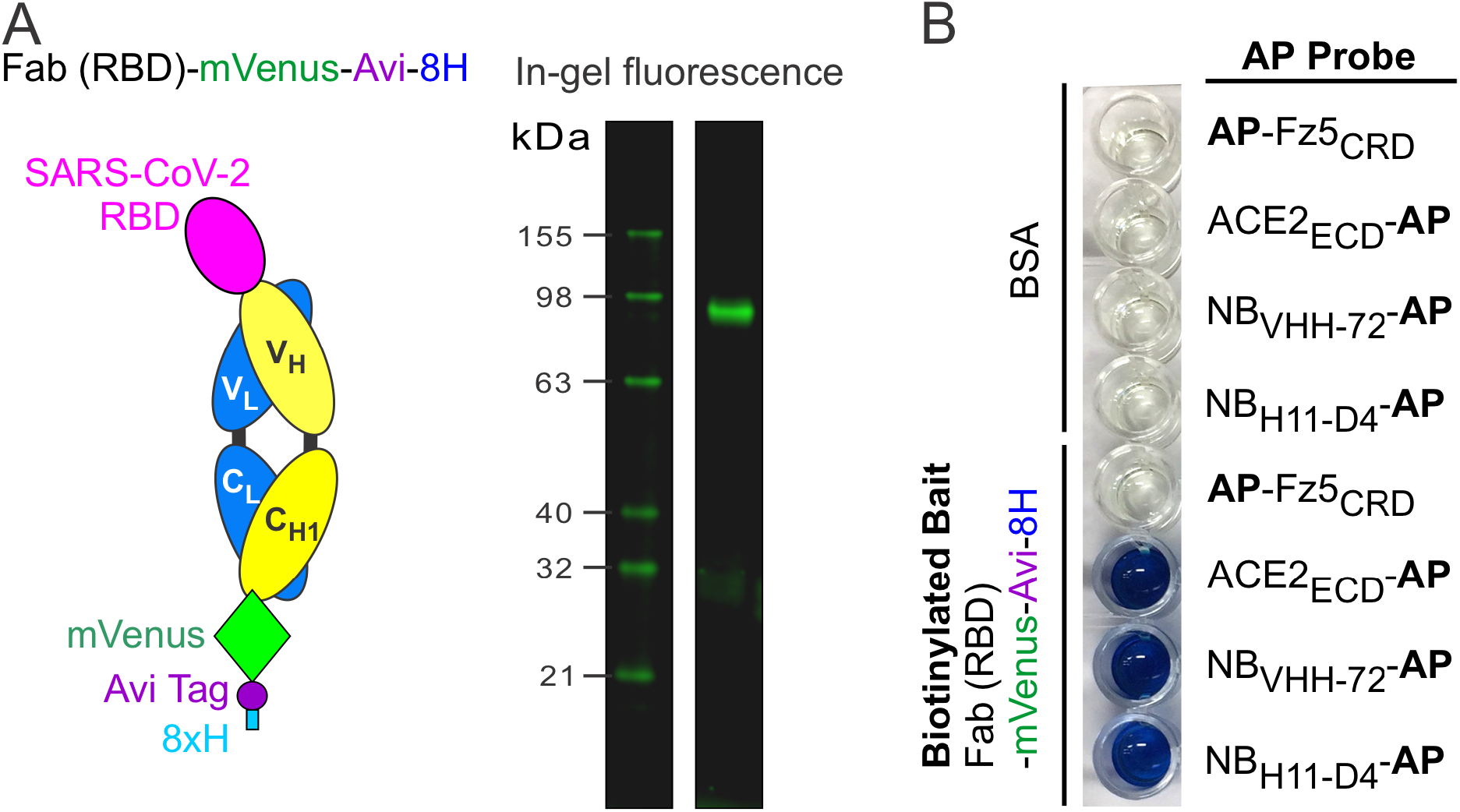
Protein expression of RBD-AD in a Fab format and AP based binding assay. **(A)** The in-gel fluorescence imaging shows the biotinylated Fab (RBD)-mVenus-Avi-8H bait that was purified by IMAC from the conditioned media and used for the binding assay. **(B)** Biotinylated Fab (RBD) baits were immobilized on streptavidin-coated wells. Bound AP fusion probes were visualized by a colorimetric AP reaction (blue).

